# Structure-Altering Mutations of the SARS-CoV-2 Frame Shifting RNA Element

**DOI:** 10.1101/2020.08.28.271965

**Authors:** T. Schlick, Q. Zhu, S. Jain, S. Yan

## Abstract

With the rapid rate of Covid-19 infections and deaths, treatments and cures besides hand washing, social distancing, masks, isolation, and quarantines are urgently needed. The treatments and vaccines rely on the basic biophysics of the complex viral apparatus. While proteins are serving as main drug and vaccine targets, therapeutic approaches targeting the 30,000 nucleotide RNA viral genome form important complementary approaches. Indeed, the high conservation of the viral genome, its close evolutionary relationship to other viruses, and the rise of gene editing and RNA-based vaccines all argue for a focus on the RNA agent itself. One of the key steps in the viral replication cycle inside host cells is the ribosomal frameshifting required for translation of overlapping open reading frames. The frameshifting element (FSE), one of three highly conserved regions of coronaviruses, includes an RNA pseudoknot considered essential for this ribosomal switching. In this work, we apply our graph-theory-based framework for representing RNA secondary structures, “RAG” (RNA-As Graphs), to alter key structural features of the FSE of the SARS-CoV-2 virus. Specifically, using RAG machinery of genetic algorithms for inverse folding adapted for RNA structures with pseudoknots, we computationally predict minimal mutations that destroy a structurally-important stem and/or the pseudoknot of the FSE, potentially dismantling the virus against translation of the polyproteins. Additionally, our microsecond molecular dynamics simulations of mutant structures indicate relatively stable secondary structures. These findings not only advance our computational design of RNAs containing pseudoknots; they pinpoint to key residues of the SARS-CoV-2 virus as targets for anti-viral drugs and gene editing approaches.

**SIGNIFICANCE:** Since the outbreak of Covid-19, numerous projects were launched to discover drugs and vaccines. Compared to protein-focused approaches, targeting the RNA genome, especially highly conserved crucial regions, can destruct the virus life cycle more fundamentally and avoid problems of viral mutations. We choose to target the small frame-shifting element (FSE) embedded in the Open Reading Frame 1a,b of SARS-CoV-2. This FSE is essential for translating overlapping reading frames and thus controlling the viral protein synthesis pathway. By applying graph-theory-based computational algorithms, we identify structurally crucial residues in the FSE as potential targets for anti-viral drugs and gene editing.

## 1 INTRODUCTION

The novel coronavirus SARS-CoV-2, the agent of the Covid-19 pandemic that has upended our world in a very short time, poses an international health emergency of unprecedented proportions. With rapid transmission and high death rates, this virus threatens the lives and livelihood of billions of people worldwide. Scientists have been quick to rise to the task to investigate collaboratively many viable paths of defense. These include development of better techniques for detection and tracing of infection, identification of viable compounds that mitigate the virus’s harm, and development of vaccines.

Some clues into the complex virus apparatus involved in the infection process are available from highly related cousins of this pathogenic coronavirus. From the SARS (SARS-CoV) and MERS (MERS-CoV) viral outbreaks, we have a reasonable mechanistic understanding of the SARS-CoV-2 virus life cycle and the infection process. This includes the cellular invasion by the viral spike (or S) protein and its binding to the ACE2 cellular receptors, and the triad of main proteins essential for producing a complete viral particle (envelope, membrane, and nucleocapsid) (1–3). In addition, scientific advances since the SARS and MERS outbreaks, along with accumulating data about the disease since its emergence in late 2019 from China, provide insights into the virology, pathology, and empirical treatments of such infections (1–3).

From this information and knowledge of other viruses like HIV-AIDS, Ebola, Influenza, Hepatitis C, or Malaria (4, 5), we have suites of anti-viral and other compounds that can potentially mitigate the damage of the virus on the body, including Remdesivir, Tocilizumab, Favipiravir, Galidesivir, Brilacidin, and Dexamethasone (5). In particular, Remdesivir, which resembles the structure of an HIV reverse-transcriptase inhibitor, has shown promise for reducing recovery time of patients in acute stages, and the steroid Dexamethasone, used widely to fight serious inflammatory reactions, was shown to reduce Covid-19 deaths. Drugs that block spike, membrane, or envelope proteins on the viral shell of coronaviruses are actively being sought, and some are being investigated in clinical trials.

With rapid deaths, as well as second and third waves of infection already seen in some countries, scientific data are critically needed to help guide clinical efforts and offer genomic, bioinformatic, and biophysical insights into effective disease treatment approaches, *both short and long term*. Because a vaccine is not imminent and new coronavirus epidemics can be anticipated, a better understanding of the viral machinery and potential therapeutic avenues that are not currently widely explored are warranted.

SARS-CoV-2, like other coronaviruses, has a ≈ 30,000 base-pair RNA genome. The viral RNA agent hijacks the host cells’ ribosome machinery to replicate and assemble itself by synthesizing a suite of viral proteins and thus continue to invade host organs. An attractive alternative line of research to protein-targeting approaches involves compounds that degrade and thus disarm the main workhorse of the virus, the RNA genome (6, 7). RNA-targeting approaches offer therapeutic potentials due to the high sequence and structure conservation of the untranslated regions of the viral genomes; they nonetheless pose physiochemical challenges due to their more complex structures. With rapid emergence of CRISPR technology, however, such approaches are becoming more viable and valuable. Indeed, CRISPR/Cas13d technology may be used in connection with a guided-RNA delivery system to edit the SARS-CoV-2 viral genome in strategic locations such as the Open Reading Frame ORF1a,b and the spike gene of the virus, with the aim of disrupting key replication or invasion processes (7). Two vaccine candidates based on the RNA are under advanced clinical trials (8) — an mRNA vaccine by Moderna, and an adenovirus vector by Johnson & Johnson.

One of the critical steps in viral replication is the translation of overlapping, but shifted, ORF1a,b gene region which codes for the unstructured polypeptides that start the cascade of viral protein synthesis. This is achieved via the process of −1 programmed ribosomal frameshifting, a mechanism to stall ribosomal translation utilized by many viruses to handle overlapping reading frames (9). The frameshifting element (FSE) is a small region (less than 100 nucleotides) in the middle of the ORF1a,b gene region; it is believed that such mechanisms rely on specific structural modules and/or associated structural transitions (10). In the SARS-CoV virus, this frame-shifting element was identified as a three-stem structure, where Stems 1 and 2 intertwine via a pseudoknot, or an intertwined base pair region (11, 12). Drug screening for this structure had suggested a potential drug to a central loop region (10, 13), and recent research suggests promise for SARS-CoV-2 as well (14).

Here we expand upon these insights for SARS-CoV-2 applications using our graph-theoretic approach, “RAG” (RNA-As-Graphs), that uses tree and dual graphs to represent, study, predict, and design RNA secondary (2D) structures (15–20). Our dual graphs represent RNA stems/helices as graph vertices and loops as edges. These graph objects can handle RNA pseudoknot motifs. By focusing on the overall connectivity/shape of the RNA 2D structural elements, RAG substantially simplifies the RNA structure, making it insensitive to the number of residues in stems and loop regions. Because our approach is general, similar techniques can be applied to other regions of the viral genome.

Here we investigate structural features of the FSE of SARS-CoV-2, and predict minimal mutations that destroy key structural features thought to be critical to its function, thus providing targets for anti-viral or gene therapy. Specifically, we use our inverse folding procedure, RAG-IF, which utilizes a genetic algorithm to systematically mutate RNA sequences to fold onto a target graph motif (21). RAG-IF relies on our computational pipeline to design RNA sequences for target graphs from constituent building blocks (22) based on fragment assembly of subgraphs (23, 24) using a library of subgraphs and corresponding 3D structures of solved RNAs (25). Such RNA design is a computationally efficient approach that can generate hundreds of candidate of sequence pools and then sort them for minimal mutations. The proposed candidates from our original design pipeline have already shown promise through experimental chemical reactivity testing (SHAPE) (24).

In this work, we first identify the SARS-CoV-2 FSE region (based on the SARS-CoV FSE sequence) and obtain its consensus structure with eight different secondary-structure prediction programs; we represent this region as dual graph 3_6, 3 stems with a pseudoknot intertwining Stems 1 and 2 (Fig. 1A-C). Next, we use RAG-IF (21), modified here to handle pseudoknots, to identify critical residues that can transform the FSE using minimal mutations to closely-related graph motifs that correspond to other known RNA structures chosen from our dual-graph motif atlas (26). Intriguingly, we find that mutating only 2-3 critical residues in the FSE can alter its structure dramatically. We also assess the stability of these mutants by compensatory mutations using the same inverse folding techniques; this essentially provides a measure of how easily ‘reversible’ these mutations are. Next, we report emerging structural insights from microsecond molecular dynamics (MD) simulations of the wildtype and mutants as additional measures of structural stability. We conclude with a summary and discussion of our findings, including a figure with key drug-target residues emerging from this work, implications to Covid-19 therapy, and future plans.

**Figure 1:**
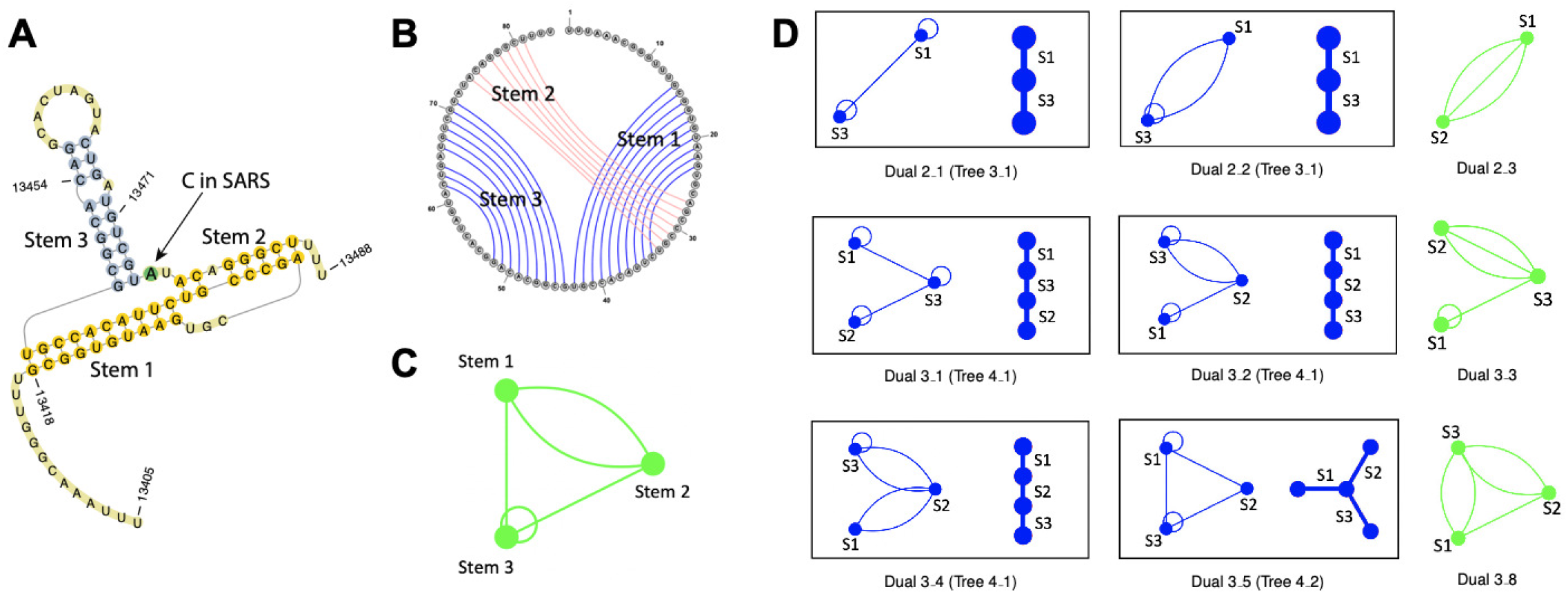
Consensus structure of the SARS-CoV-2 frame-shifting element (FSE) and related dual graph targets. (A) 2D structure of the FSE, a three-stem pseudoknot motif. (B) Circular plot of the FSE 2D structure, where the pseudoknot corresponds to the crossing of the two base pairs. (C) Associated dual graph motif 3_6 for the FSE 2D structure. (D) Dual graphs with 2 or 3 vertices related to the FSE 3_6 motif. In dual graphs, stems in RNA secondary structures are vertices, while junctions, bulges and loops are edges (15). The targets containing pseudoknots are shown in green. Those not containing pseudoknots, are also represented as tree graphs, where stems denote edges and junctions, bulges and loops are vertices.

## 2 MATERIALS AND METHODS

### 2.1 Identification of the SARS-CoV-2 FSE

The SARS-CoV FSE has been identified as the slippery site UUUAAAC, and a downstream three-stem pseudoknot-containing structure (12, 27). Taking this information as a guide, we used RNAMotif (28) to search for the same nucleotide pattern (bracketed by 5’UUUAAAC, an H-type pseudoknot, and UUU3’) in SARS-CoV-2 complete RNA genome (GenBank entry MT246482). This led to the identification of a 84-nucleotide sequence in the viral genome of SARS-CoV-2, with a single-nucleotide substitution as compared to SARS-CoV FSE (Fig. 1A).

### 2.2 Secondary Structure Prediction Packages

We use eight RNA 2D structure prediction packages that handle pseudoknots to predict the 2D structures of 84-residue and 77-residue (without the slippery site) FSE of SARS-CoV-2: PKNOTS (29), NUPACK (30), IPknot (31), Hotknot (32), Vfold2D (33), SPOT-RNA (34), ProbKnot (35), and vsFold5 (36). PKNOTS and NUPACK were installed locally, and webservers were used for the others. The structures were first predicted using the default parameter sets for all eight packages; if the default set did not yield the expected three-stem pseudoknot structure, other parameter sets available on the webservers were also utilized.

Fig. S1 and Fig. S2 show the minimum free energy structures predicted by the eight programs, using the default or alternative parameter set, for both the 84-residue and 77-residue FSE.

### 2.3 RAG Dual Graphs

In RNA 2D structures, double-stranded (base-paired) regions are called stems/helices, and single-stranded regions are called loops (15–17). The rules for representing 2D structures as dual graphs are:

1. Stems are represented as vertices and loop strands are represented as edges.
2. Stems must have at least two base pairs to be considered as vertices.
3. Each single strand in bulges, internal loops and junctions are represented as an edge that connects vertices. Bulges/internal loops with only one residue on each strand are ignored.
4. Hairpin loops are represented as self edges.
5. Unpaired residues at the 5’ and 3’ ends are ignored.

Fig. 1A,C show the 2D structure of the SARS-CoV-2 FSE and its corresponding dual graph, 3_6. Our dual graph enumeration algorithm (26) uses graph theory to describe all possible dual graphs for 2-9 vertices. There are 110,667 dual graphs in our current dual graph library, and we show those with 2-5 vertices in Fig. S3.

### 2.4 RAG-IF for Minimal Mutations

We use our inverse-folding protocol RAG-IF (21) to mutate the 77-residue FSE sequence to fold onto closely related dual graph motifs. Our RAG-IF program defines the inverse-folding with mutations section of our computational pipeline for novel RNA design (21, 24). For a target tree graph (i.e., no pseudoknots), our original design pipeline generates a pool of RNA sequences, some of which fold onto the target graph, as determined by two 2D structure prediction packages. For the unsuccessful ones, RAG-IF is applied to determine minimal mutations that lead these sequence to fold onto the target topology.

RAG-IF has three steps: (1) identify mutation regions and target 2D structures, (2) produce candidate sequences by mutations by a genetic algorithm (GA), and (3) optimize the mutated sequence pool by sorting and retaining only minimal or essential mutations that fold onto the target graph.

For tree graphs, all three steps are automated. However, to use RAG-IF for dual graphs capable of handling pseudoknots, we manually choose the target residues for mutation (step 1), modify aspects of the GA for speed, and change the 2D structure prediction packages to ones that can handle pseudoknots for steps 2 and 3.

#### 2.4.1 Selection of mutation regions

We compare the wildtype FSE graph and the target graph to identify the smallest possible mutation regions required for the transformation. Specifically, we focus on breaking the FSE pseudoknot/Stem 2. Here we describe how this is accomplished for our four most successful target graphs 3_5, 3_2, 2_1, and 3_3 (shown in Fig. 1D and Fig. S4).

Target dual graph 3_5 is a three-way junction, without any pseudoknots, and it corresponds to RNAs like the thi-box riboswitch. Two of the three stems of the FSE (Stems 1 and 3) are also present in the 3_5 graph, hence are kept intact in the transformation. To break the FSE pseudoknot/Stem 2, we seek residues at the 3’ end of the FSE to form a new stem with the 5’ end. Hence, we select residues 71-77, part of Stem 2 and the 3’ end, as targets for mutation.

Target graph 3_2 has 3 stems connected by a single-stranded region and an internal loop, and it includes the original Stem 3. This graph corresponds to RNAs like U6 snRNA. For this transformation, we use the 5’ end and hairpin loop of Stem 1 (Fig. S4) to form a new stem. To pair the residues involved, we select the 3’ strand of the original Stem 1, residues 27-33, as the mutation region.

Target graph 2_1, corresponding to RNAs like the cleaved hammerhead ribozyme, has two stems and a connecting single-stranded region. For this transformation, we aim to keep Stems 1 and 3 intact, but destroy Stem 2, while avoiding two other potential pseudoknots (shown in Fig. S4). Hence, we select residues 22-24 and 70-74 of Stem 2 and the 3’ end residues 75-77 for mutation.

Target graph 3_3 has 3 stems and a structurally different pseudoknot. It corresponds to RNAs like the IRES (PDB ID: 2IL9). To break the FSE pseudoknot, we want the 3’ strand of Stem 2 to bind with the hairpin loop of Stem 3 to form a new pseudoknot. However, to avoid a another potential pseudoknot as shown in Fig. S4, we select residues 22-24 and the 3’ strand of Stem 2 (residues 68-73) as mutation regions.

#### 2.4.2 Genetic algorithm application

Genetic algorithms (GA) mimic evolution in Nature. A population with a defined *fitness* undergoes random mutation, crossover and selection, and those with high fitness are retained. In our case, the individuals in the population are RNA sequences, and the fitness is the number of residues that have the same 2D structure in both the target and the predicted structure. Based on prediction performance on the 77-residue FSE and computational complexity (see below), we choose NUPACK to be the prediction program used in the GA. The initial population is obtained by randomly assigning nucleotides to the mutation region in the RNA sequence. This population is then subject to 100 iterations of random mutation, crossover, selection and nomination (see (21) for details).

For minimal mutation search, we use a population of 100 sequences (and 10 sequence replacement in the selection step). The GA terminates if 200 high-fitness sequences are nominated or the execution time exceeds 6 hours. The nominated sequences are further screened by prediction program PKNOTS, and only those that satisfactorily fold onto the target graph by both NUPACK and PKNOTS are retained.

For compensatory mutation search, the starting sequences are the mutants, the mutation regions are the same, and the target folding is the 2D structure predicted by NUPACK for the wildtype 77-residue FSE. Expanded RAG-IF runs (500 sequences) are taken for targets 2_1 and 3_3, with 50 sequence replacement instead of 10.

#### 2.4.3 Sequence pool optimization

During mutation optimization, we remove any mutations that do not change the target fold (dual graph) of the sequence. The remaining mutations are deemed as essential or minimal (21).

### 2.5 Molecular Dynamics Simulation Details

The 3D structures of the 77-residue wildtype FSE, along with three mutants, 72C-74C, 30G-32C, 23G-73A, were predicted using 4 structure prediction webservers: RNAComposer (37), Vfold3D (38), SimRNA (39), and 3dRNA(40). We discard any structure that did not correspond to the intended dual graph motif (3_6 for the wildtype, 3_5 for 72C-74C mutant, 3_2 for 30G-32C mutant, and 2_1 for 23G-73A mutant) based on the 2D structures annotated by either DSSR (41) or RNAView (42). Structure validation was performed by MolProbity (43) (using nuclear hydrogen positions), and the structure with least steric clashes and best geometry was selected for the subsequent MD simulations. For the wildtype as well as the 3 mutants, the selected structure was predicted by RNAComposer.

MD simulations were performed using Gromacs 2020.3 (44), with the Amber OL3 forcefield (45). The systems were solvated with TIP3P water molecules in the cubic box whose boundaries extended at least 10 Å from any RNA atom (46). Sodium ions were randomly placed for charge neutralization, and additional Na^+^ and Cl^−^ were added for 0.1M bulk concentration. The total number of atoms for the simulations of the wildtype, 72C-74C mutant, 30G-32C mutant, and 23G-73A mutant are 119671, 156619, 162175, and 151076, respectively.

The systems were energy minimized via steepest descent and equilibrated while fixing the RNA. Simulations were run with a timestep of 2 fs and a SHAKE-like LINCS algorithm (47) with constraints on all bonds. The equilibration was performed for 100 ps in the NVT ensemble (300 K) and then 100 ps in NPT ensemble (300 K and 1 bar). Production runs were performed for 1 *µ*s under NPT. The RNA and ionic solvent were independently coupled to external heat baths with a relaxation time of 0.1 ps. The Particle Mesh Ewald method (48) was used to treat long-range electrostatics. The short-range electrostatic cutoff is set at 10 Å, and the Van der Waals cutoff is set at 10 Å.

The trajectories were analyzed using Gromacs, and structure visualization was prepared by PyMOL (49) and VMD (50). Root mean square deviations (RMSD) and root mean square fluctuations (RMSF) were calculated with reference to the NPT equilibrated structure using all heavy RNA atoms. Cluster analysis was conducted via Gromos method with the RMSD cutoff of 3.5 Å on RNA non-H backbone atoms. Principal component analysis (PCA) was performed using all heavy atoms in the RNA. Structures used for clustering and PCA analyses were taken every 250 ps from 500ns to 1 *µ*s.

## 3 RESULTS

### 3.1 Secondary Structure and Dual Graph for SARS-CoV-2 FSE

Based on genomic sequencing, SARS-CoV-2 genome is 89% similar to SARS-CoV and 50% to MERS-CoV. From the study of earlier RNA viruses, the FSE consists of two components: a heptamer of the form XXXYYYZ, known as the slippery site, where the tRNA dissociates from the mRNA and the ribosome ‘slips’ to an overlapping reading frame; and a downstream mRNA element, usually a pseudoknot (9, 51). Specifically, the SARS-CoV FSE has been identified as the slippery site UUUAAAC, and a downstream three-stem pseudoknot-containing structure (12, 27). Because of strong sequence conservation, it is presumed that this similar FSE motif would also exist in SARS-CoV-2. We searched for this FSE motif (see Identification of the SARS-CoV-2 FSE) and identified the same 84-nucleotide sequence in the viral genome of SARS-CoV-2 (residues 13405–13488, analogous to the SARS-CoV residues 13392–13475), with a mere single-nucleotide substitution (A in SARS-CoV-2 instead of C in SARS-CoV in position 72 of this sequence), as shown in Fig. 1A.

To assess the secondary (2D) structure of the SARS-CoV-2 FSE and to select software for our mutational analysis, we applied eight 2D-structure prediction packages that can handle pseudoknots. See Secondary Structure Prediction Packages for computational details.

Out of these eight packages, 3 programs predicted a three-stem pseudoknot structure (Fig. S1). As the remaining programs predicted additional stems or pseudoknots that involved the heptamer slippery sequence, we applied the 8 programs again to the 77-residue system without the slippery site. Now 5 of the 8 programs predict a three-stem pseudoknot structure (Fig. S2).

Our resulting consensus three-stemmed pseudoknot structure in Fig. 1A shows the stem regions that lead to an intertwined structure in three dimensional space. The corresponding circular plot shows the base pairing. Using our rules to represent RNA 2D structure for RAG dual graphs (RAG Dual Graphs), this FSE structure corresponds to the dual graph 3_6 from our recently enumerated atlas of dual graphs up to 9 vertices (26). The three stems correspond to three vertices, and the loops correspond to connecting edges. This 3_6 fold motif also corresponds to multiple riboswitch structures of SAM (PDB IDs: 3NPN, 3NPQ) and flouride riboswitches (PDB IDs: 3VRS, 4EN5, 4ENA, 4ENB, and 4ENC) (26).

Although it is not conclusively known what the FSE structure is in SARS-CoV-2 and whether this region’s folding may be affected by neighboring sequences and its binding to the nucleocapsid protein (see Conclusion and Discussion), other recent works have used various computational tools and experimental input to suggest its structure (14, 52).

Since our dual graphs are independent of the number of residues in stems and loops, the graph motif 3_6 corresponds to every three-stem pseudoknot structure predicted by the above programs for both the 84-residue and 77-residue FSE. Thus, the insensitivity of our graph-based approach to small differences in base pairing annotations makes its application more robust. We use the 77-residue FSE and software packages NUPACK and PKNOTS for our computational mutational analysis described here.

### 3.2 Identification of Minimal Mutations To Destroy Stem 2 and/or Pseudoknot

The SARS-CoV-2 FSE structure and associated transitions are thought to be key for ribosomal frameshifting and the subsequent translation of the ORF1a,b region of the viral genome. Any disruption of key structural features would possibly inhibit or hamper this translation. Therefore, we seek crucial FSE nucleotides, especially those that destroy Stem 2 (Fig. 1) and the pseudoknot, that can potentially serve as antiviral drug targets.

Identifying these crucial residues is challenging, even in a relatively small sized RNA like the FSE. Fortunately, our RAG representations allow us to simplify this problem. We formulate our goal here as identifying minimal mutations to transform the 3_6 dual graph motif of the FSE into nine other dual graphs, of equal or smaller size, that correspond to 2D structures of known RNAs (26). These nine targets are shown in Fig. 1D. Among these, 6 graphs are pseudoknot free (blue) and the remaining 3 contain pseudoknots (green).

We use our inverse-folding protocol RAG-IF (21) to mutate the FSE sequence, as described in Materials and Methods. After the minimal mutations are identified, we transform the top mutated sequences back to the original FSE graph 3_6 using RAF-IF to access the mutants’ stability. We want to determine whether we recover the original FSE sequence and whether other compensatory mutations can accomplish structural transformations.

Below, we describe minimal mutation and compensatory mutation results for four of the most successful targets, 3_5, 3_2, 2_1, and 3_3, in turn (see Fig. 2, 3, 4, 5). Summary Fig. 6A shows residues involved.

**Figure 2:**
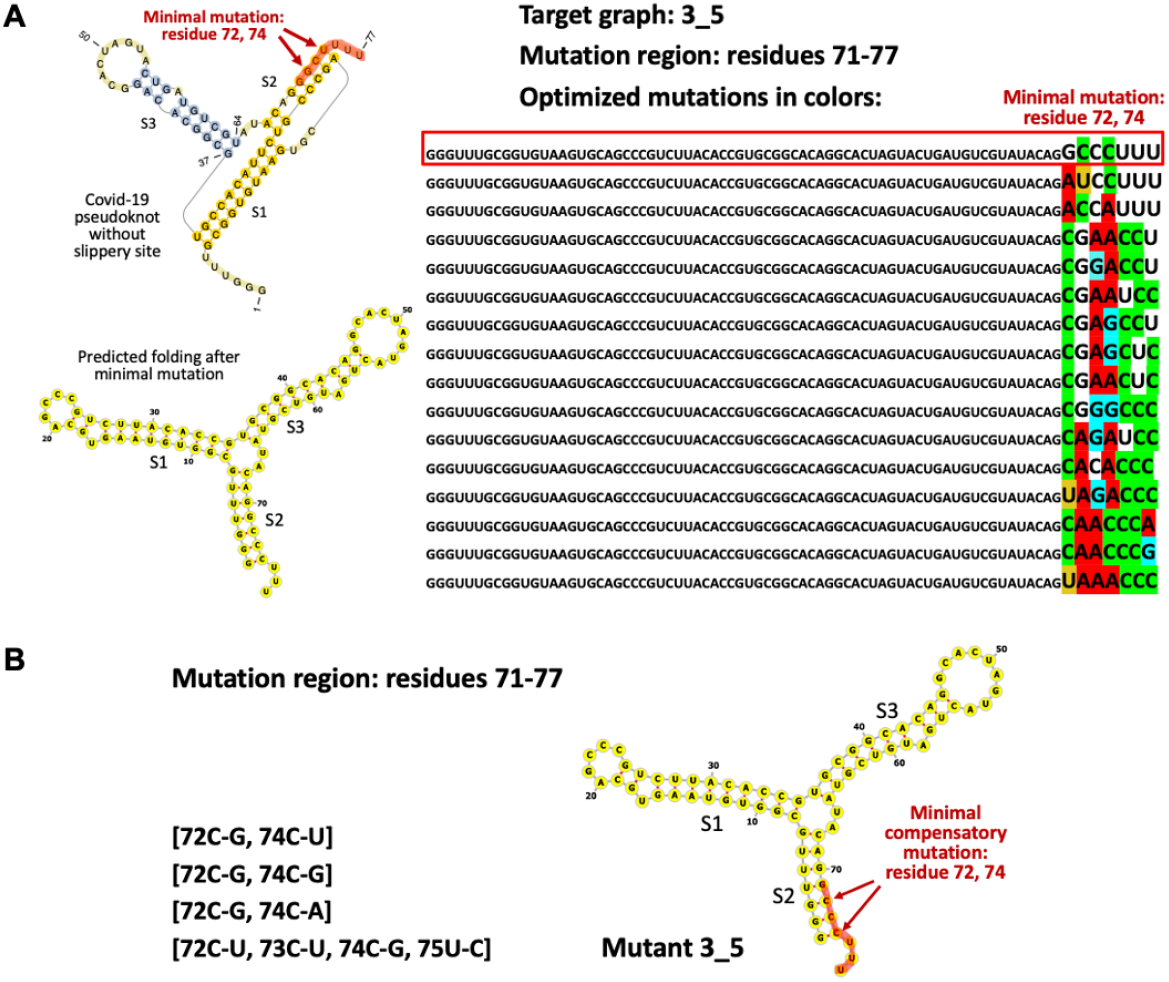
Mutation results for the 3-way junction target graph 3_5. (A) Minimal mutations: mutation candidate region is highlighted in red. Point mutations are colored according to the nucleotides. One top mutation is selected, whose predicted secondary structure is shown. (B) Minimal compensatory mutations.

**Figure 3:**
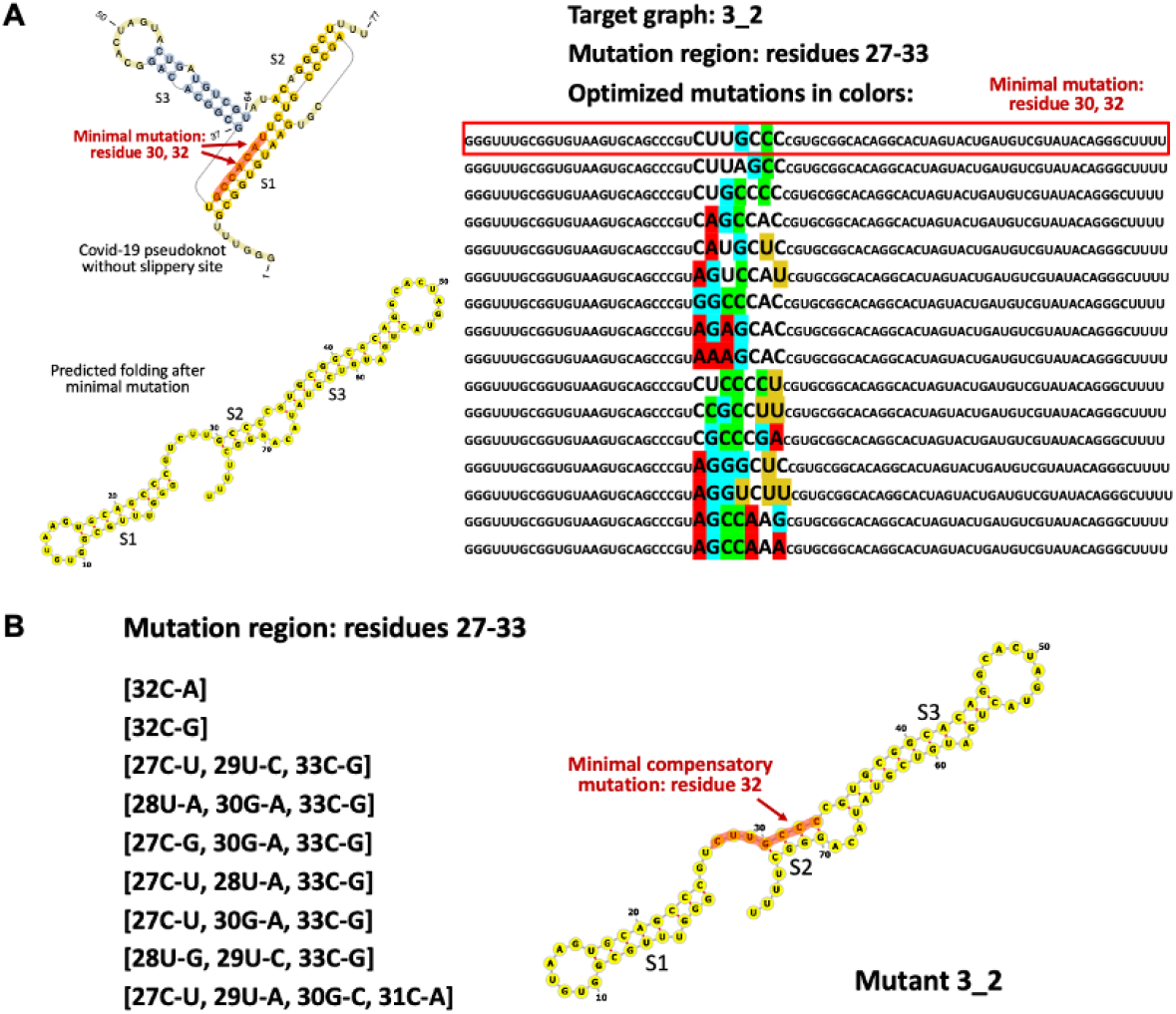
Mutation results for the 3-stem with internal loop target graph 3_2. See Fig. 2 caption.

**Figure 4:**
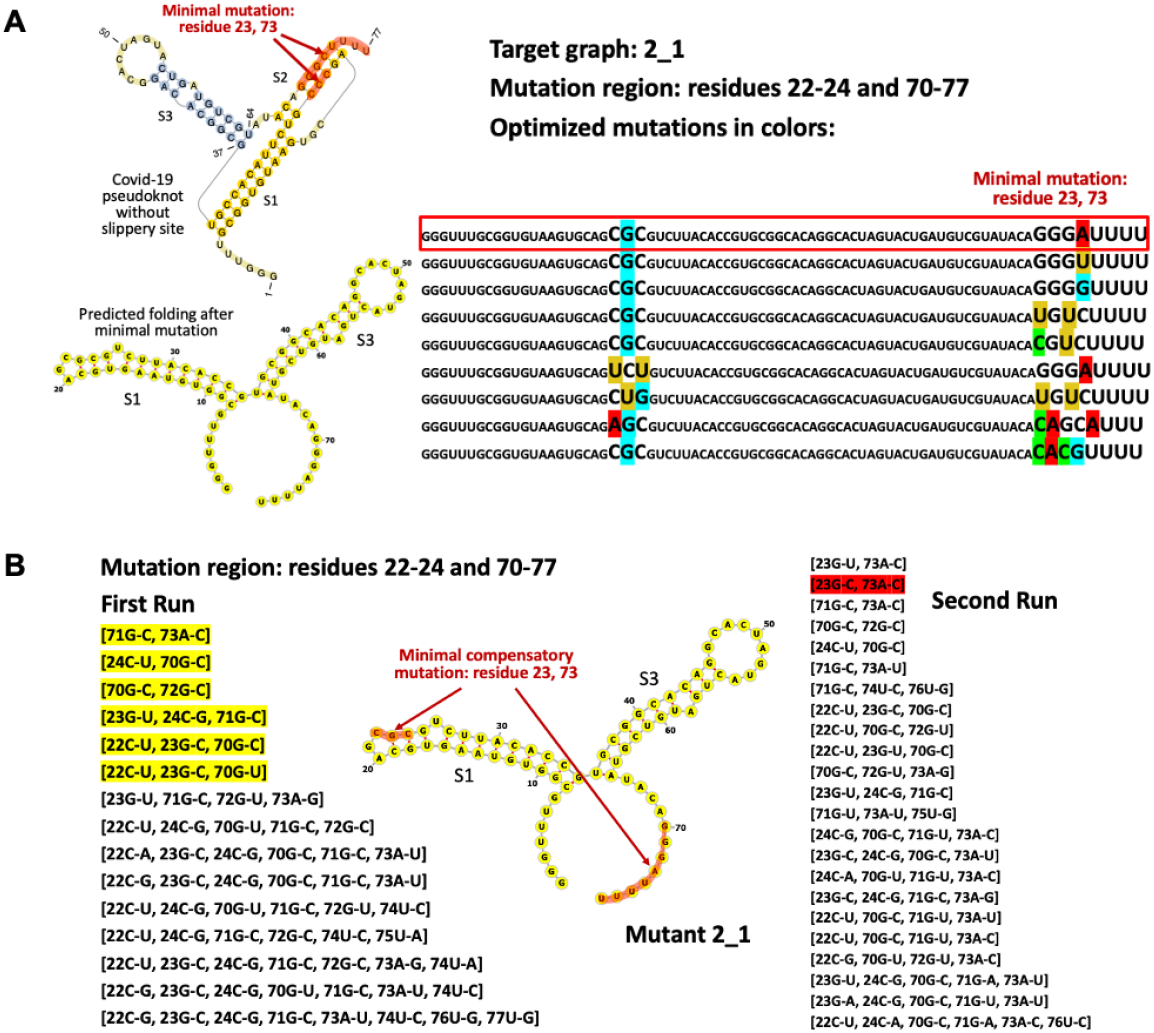
Mutation results for the 2-stem target graph 2_1. See Fig. 2 caption.

**Figure 5:**
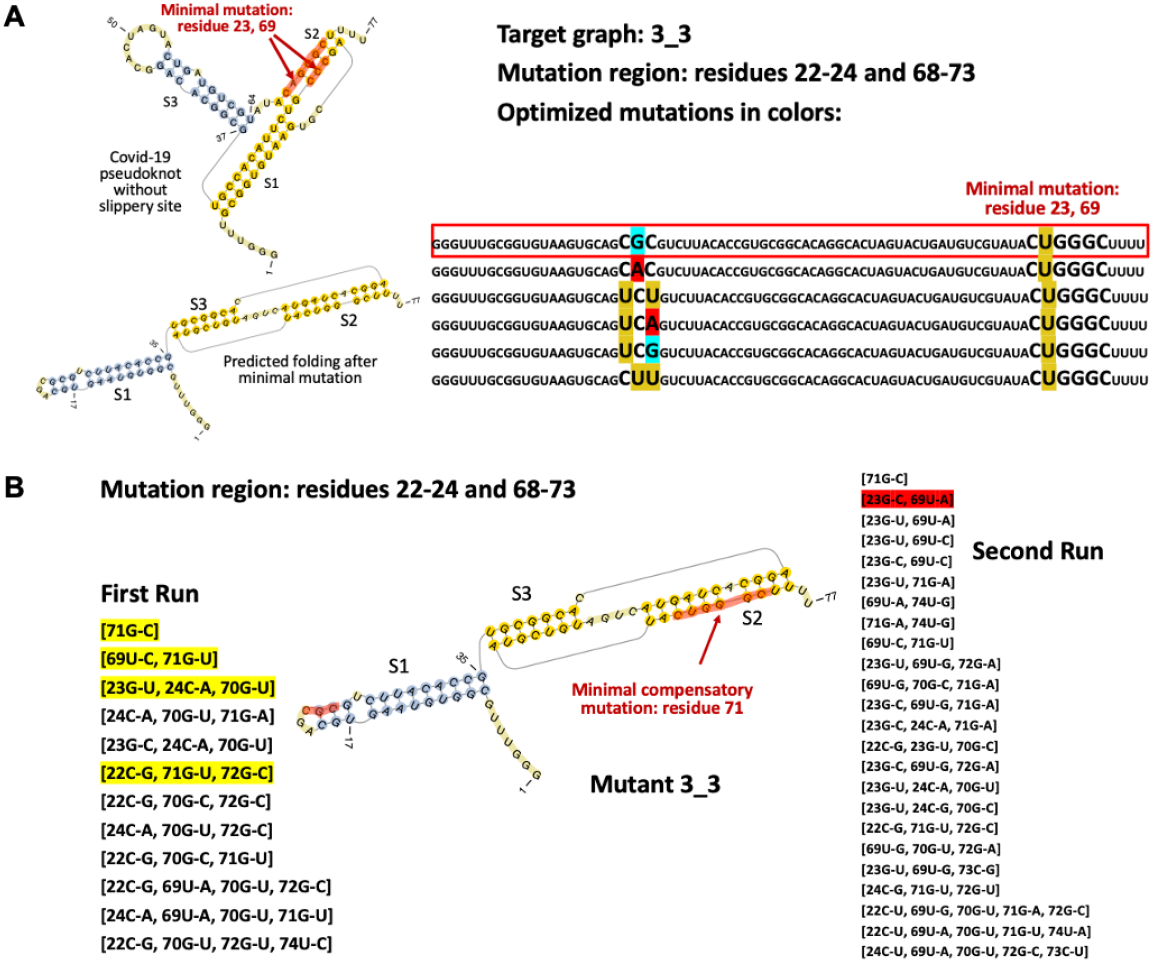
Mutation results for the 3-stem with different pseudoknot target graph 3_3. See Fig. 2 caption.

**Figure 6:**
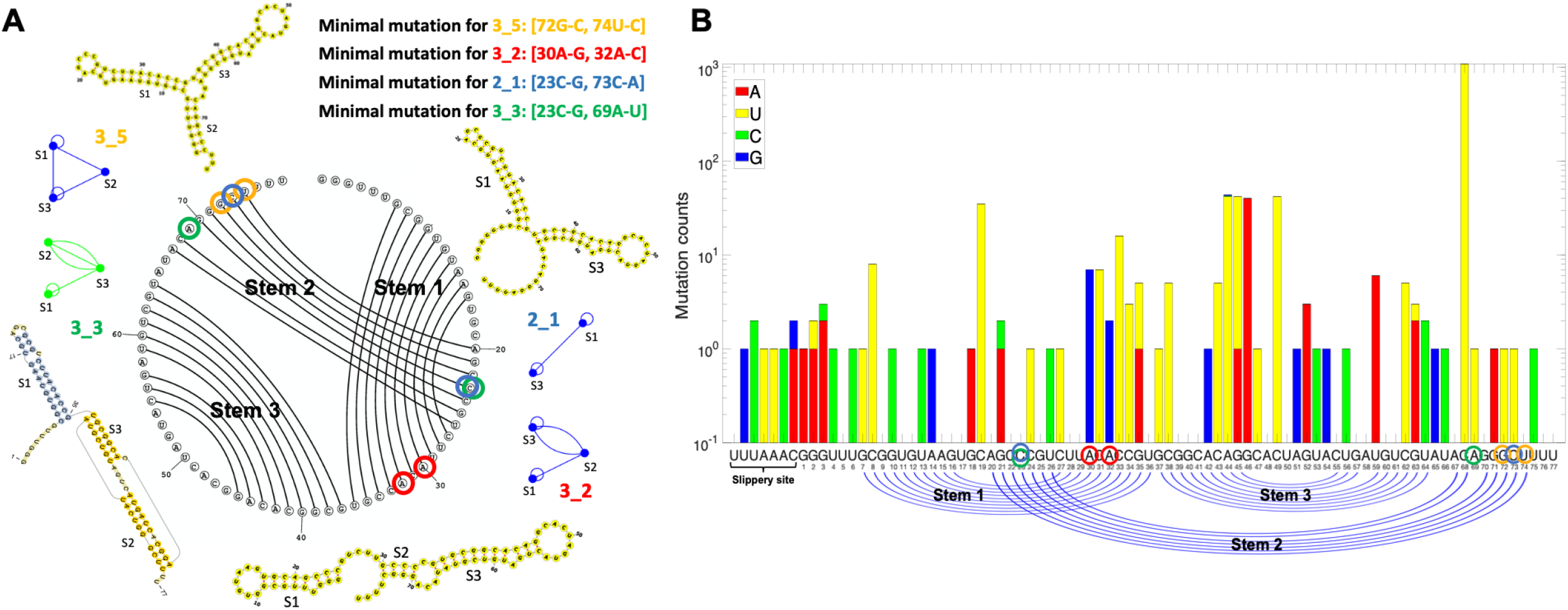
FSE residue analyses. (A) Minimal mutation residues are circled for the different mutants: yellow for 3_5, red for 3_2, blue for 2_1, and green for 3_3. (B) FSE mutation findings for Covid-19 variants from the GISAID database (53) as of August 10, 2020.

#### 3.2.1 Mutations for target graph 3_5 (3-way junction)

To transform 3_6 of the FSE into target 3_5 (Fig. 2A), we keep Stem 1 and 3 intact and select residues 71-77 (highlighted in red) in Stem 2 for mutation.

RAG-IF GA produces 115 sequences that fold onto the target graph 3_5 by both NUPACK and PKNOTS, with number of mutations ranging from 3−7 (Table S1). Following mutation optimization to remove non-essential mutations, we obtain 16 unique sequences with 2−7 minimal mutations. Significantly, [72G-C, 74U-C] destroys the corresponding base pairs in Stem 2 of the FSE and strengthens the new stem of 3_5 with G-C base pairs.

The same outcome for this transformation of 3_6 to 3_5 using [72G-C, 74U-C] comes from two additional prediction programs (see Fig. S5). Thus, we consider this the top mutant choice for target 3_5. Its predicted 2D structure is shown in Fig. 2A.

Transforming this mutant [72G-C, 74U-C] back to the 3_6 graph can be accomplished by 2 double mutants besides [72C-G, 74C-U]: [72C-G, 74C-G] and [72C-G, 74C-A] (Fig. 2B).

#### 3.2.2 Mutations for target graph 3_2 (3-stem with internal loop)

Next, we transform the FSE into target graph 3_2 (Fig. 3A). To construct 3_2 from 3_6, we keep Stem 3 intact, and select the mutation region as residues 27-33.

RAG-IF GA generates 64 candidate sequences that fold onto the target 3_2 by both NUPACK and PKNOTS, with mutations ranging from 3−7 (see Table S1). Following mutation optimization, we obtain 16 unique sequences, with 2−6 minimal mutations. Two of the 16 optimized sequences require only 2 mutations: either residues 30 (from A to G) or 31 (from C to G), with residue 32 (from A to C).

Both sequences mutate key residues to destroy Stem 1 of the FSE, along with the hairpin loop that forms the pseudoknot (Stem 2), and strengthen the newly formed stems with G-C base pairs. The predicted structure for mutation [30A-G, 32A-C] (Fig. 3A) is confirmed by another prediction program (Fig. S5).

When transforming the mutant [30A-G, 32A-C] back to the 3_6 graph, we obtain 55 candidate sequences before optimization from the GA. Besides recovering [30G-A, 32C-A], we find two other single-point mutations, [32C-A] and [32C-G], that recover the graph 3_6. In fact, 48 of the 55 sequences mutate residue 32. Following optimization, we obtain 9 unique sequences (Fig. 3B).

Interestingly, most of the mutations from 44 of the 55 candidates are deemed non-essential, and only [32C-A] or [32C-G] are retained. This includes the reverse mutation [30G-A, 32C-A]. Mutating residue 32 breaks the middle base pair in a relatively small stem in the 3_2 mutant, thus destabilizing the whole structure. If residue 32 is not mutated, at least 3 minimal mutations are required, rendering residue 32 as one of the key residues for structural transformations.

#### 3.2.3 Mutations for target graph 2_1 (two stems)

To transform 3_6 of the FSE into 2_1, we aim to destroy Stem 2, and select residues 22-24 and 70-77 as mutation candidate regions.

RAG-IF GA produces 221 candidate sequences that fold onto the target graph 2_1 by both NUPACK and PKNOTS: the number of mutations ranges from 6−11 (Table S1). After mutation optimization, we retain 9 unique sequences (shown in Fig. 4A) with 2−5 minimal mutations. The three sequences with only 2 mutations involve residues 23 (from C to G) and 73 (from C to A, G, or U). Because Stem 2 in the FSE is mainly formed by four G-C base pairs, mutating 23 and 73 breaks two of them which destabilizes the stem.

The mutant RNA [23C-G, 73C-A] sequence also folds onto our target graph 2_1 by two additional prediction programs (Fig. S5). The predicted 2D structure, shown in Fig. 4A, has Stem 1 longer than that of the original FSE.

Our reverse transformation of the mutant [23C-G, 73C-A] to 3_6 generates 23 unique sequences after an expanded pool (see Materials and Methods), recovering [23G-C, 73A-C] and finding many others (Fig. 4B).

#### 3.2.4 Mutations for target graph 3_3 (3-stem with different pseudoknot)

To transform 3_6 of the FSE into 3_3 (Fig. 5A), we destroy Stem 2 to form the new pseudoknot, by mutating residues 22-24 and 68-73, the two strands of Stem 2.

RAG-IF GA produces 200 candidate sequences that fold onto the target 3_3 by both NUPACK and PKNOTS, with mutations ranging from 4−9. Following optimization, 6 unique sequences emerge with only 2−3 minimal mutations. Two sequences that require two mutations, involve residues 23 (from C to G or A) and 69 (from A to U).

Interestingly, all unique sequences require mutating residue 69 to U, which is a bulge in the FSE Stem 2. Although mutating residue 69 to U creates an A-U base pair for the new pseudoknot, residue 69 likely has a more important stabilizing role for the original Stem 2. The RNA sequence with mutation [23C-G, 69A-U] in Fig. 5A is supported by another prediction program (Fig. S5).

Reverse transformation of mutant [23C-G, 69A-U] onto graph 3_6 with a large pool of sequences generates 24 unique sequences, 8 of which require only 2 mutations. Four of these sequences mutate the same residues 23 and 69, including the mutation that recovers the wildtype FSE [23G-C, 69U-A] (highlighted in red).

In addition, one single-point mutation [71G-C] is also found. Residue 71 is not one of the original mutation residues, but it is base paired with residue 23 in the wildtype FSE pseudoknot. Therefore, the compensatory mutation [71G-C] restores this G-C base pair and hence the original pseudoknot.

#### 3.2.5 Summary of minimal mutations

Transformations of the FSE into our four target RNAs utilize unpaired residues to form new stems, often keeping Stem 3 intact, and focusing on Stem 2 (Fig. 6A). Residue 23 is mutated from C to G in the top minimal mutation for both 2_1 and 3_3. This mutation breaks the G-C base pair in the original Stem 2 and elongates Stem 1 by forming new G-C base pair.

In the reverse transformations back to 3_6, the original mutations are recovered by RAG-IF, and single mutations are also found in the case of 3_2 and 3_3. Because RAG-IF aims to recover the original graph 3_6, not the original FSE sequence, compensatory mutations that require fewer residues than the forward mutations are possible. In 3_5, all compensatory mutations require the original 2 residues, so we consider this design the most stable one.

For other target graphs shown in Fig. 1D, such as 3_1 and 3_8, at least 4 residues are required for minimal mutations, and for graph 2_2, 14 residues are required (see Table S2).

### 3.3 Molecular Dynamics Analysis of Wildtype and Mutant Systems

To assess the stability of the wildtype FSE of SARS-CoV-2 and the top 3 mutants predicted by mutational analysis (3_5 [72G-C, 74U-C] mutant, 3_2 [30A-G, 32A-C] mutant, and 2_1 [23C-G, 73C-A] mutant), we subject the solvated systems to MD simulations of 1 *µ*s. To develop initial tertiary structures for these systems, we use four 3D structure prediction webservers: RNAComposer (37), Vfold3D (38), SimRNA (39), and 3dRNA (40).We select one structure each for the wildtype and the 3 mutants for MD simulation (Materials and Methods).

RMSD over the 1 *µ*s simulations is a measure of the conformational stability of the predicted structures (Fig. 7). Since predicted (and not experimentally determined) 3D structures are used for the simulations, high RMSD values (in 10s Å here) with reference to the starting structures is expected. However, the simulation settles down within the first 250-300 ns (with the 3_2 mutant taking the longest), indicating that the wildtype and the 3 mutant systems have reached a relatively stable state. The residues that fluctuate most are the unpaired residues in the loop regions, as indicated by the RMSF plots in Fig. 7. The 3_2 mutant is again the exception here, likely due to the time taken to reach relative stability.

**Figure 7:**
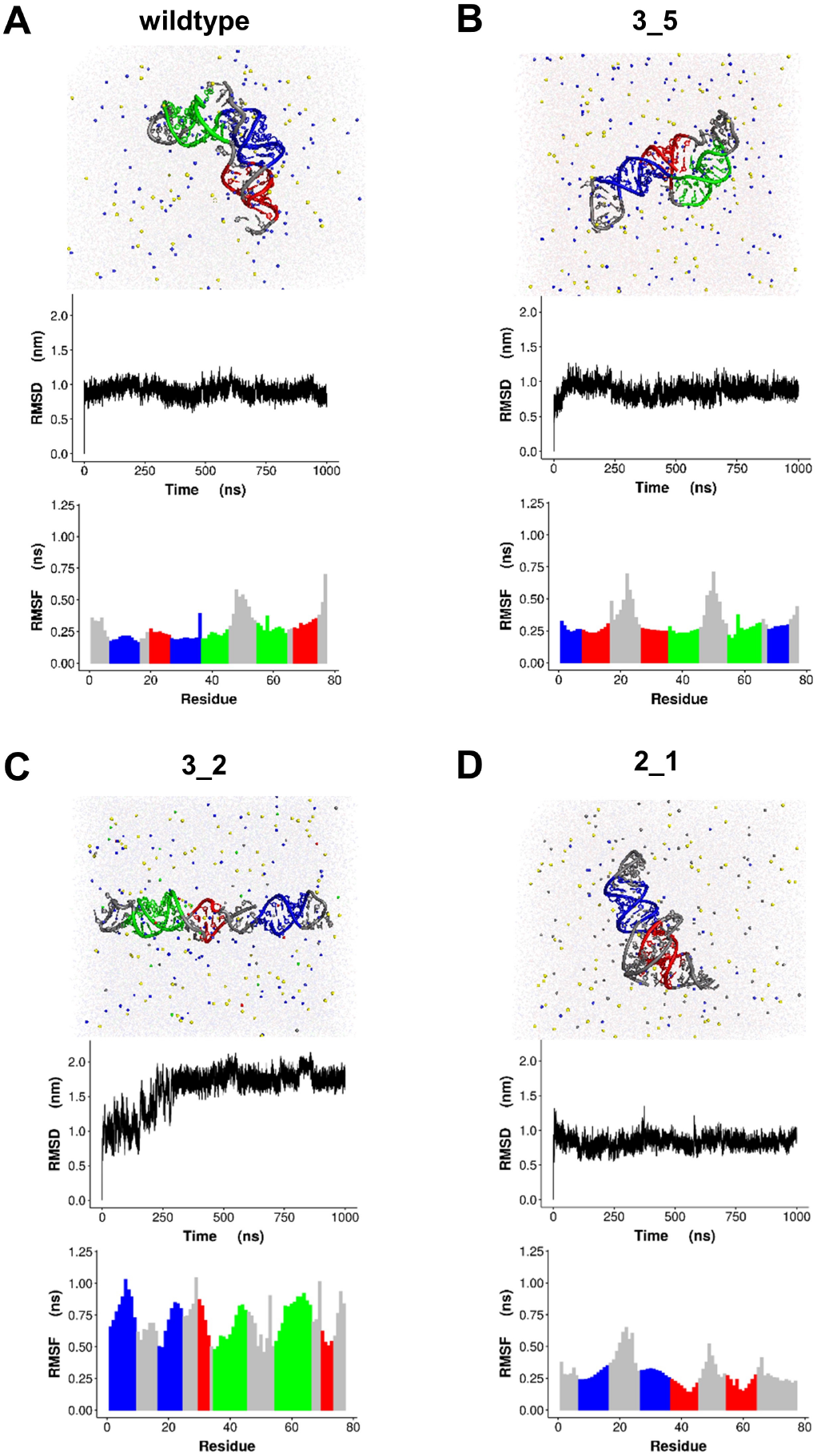
MD simulations of the wildtype and selected mutant systems using predicted models as initial structures. Snapshots of the wildtype and mutant systems at 1 *µ*s along with RMSD and RMSF are presented for (A) wildtype, (B) 3_5 mutant, (C) 3_2 mutant and (D) 2_1 mutant. The screenshots and RMSF plots are color-coded for each RNA: (A) residues 7-16 and 27-36 in blue, residues 20-26 and 67-74 in red, residues 37-45 and 55-64 in green; (B) residues 1-7 and 68-74 in blue, residues 27-35 and 8-16 in red, residues 36-45 and 55-65 in green; (C) residues 1-9 and 17-24 in blue, residues 30-33 and 70-73 in red, residues 35-45 and 55-66 in green; (D) residues 7-16 and 27-36 in blue, residues 37-45 and 55-64 in red.

To study changes in 2D structures of the wildtype and mutants, we clustered the conformations between 500ns to 1 *µ*s (see Materials and Methods). The dominant cluster for the wildtype contains roughly 73% of the total conformations, indicating the stability of the system. The values are 50-60% for the 3_5 and 2_1 mutants, and around 23% for the 3_2 mutant, suggesting that order of increased flexibility. In addition, the representatives of all top clusters (more than 50 conformations) for the wildtype, the 3_5 and the 2_1 mutant, and all but one top cluster for the 3_2 mutant have the intended dual graph motif. This indicates that the even though small changes in 2D structures occur over the course of the simulation (see Fig. S6), our dual graph motifs and predicted mutations can lead to stable structures.

We also dissect the dominant motions of the wildtype and the 3_5 mutant with PCA (see Fig. 8). The wildtype system has global bending of loop 3 as PC1, and stem twisting for PC2 and PC3. The main motions of 3_5 mutant are bending of terminal loops and twisting of the stems. It is possible that these motions are involved in large-scale structural transitions, but more extensive simulations with possibly enhanced sampling techniques are needed.

**Figure 8:**
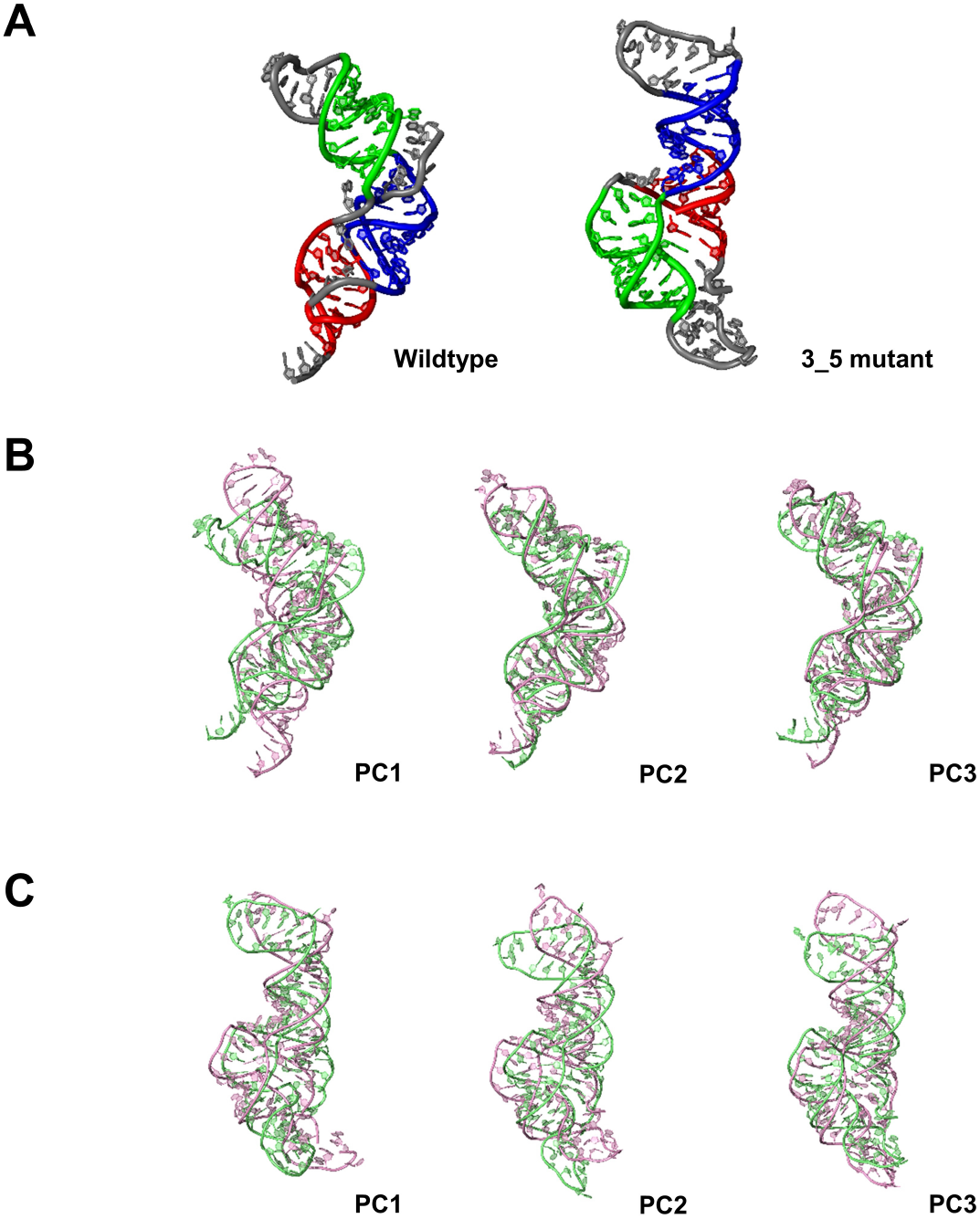
Dominant structure and motions of the wildtype and 3-way junction (3_5) mutant from MD simulations. (A) The largest cluster center structures from last 500 ns in the 1 *µ*s MD trajectories of wildtype and 3_5 mutant. (B-C) Motions of (B) the wildtype and (C) 3_5 mutant by top 3 principal components (PCs). Screenshots were extracted as two extreme structures on selected PC.

Our electrostatic surface (generated using the CHARMM-GUI PBEQ solver at http://www.charmm-gui.org/?doc=input/pbeqsolver) analysis of the wildtype system reveals some positively-charged regions along the mostly negative surface, and some grooves (Fig. 9). Some of these positively charged regions and grooves are found near residues 75-77 at the 3’ end of the FSE, residues 26-27 at the juncture of Stems 1 and 2, base pair 10-33 in Stem 1, residues 3-5 at the 5’ end of the FSE, and residues 47-53 near the hairpin loop of Stem 3. Interestingly, some of these regions coincide with the crucial residues identified by our mutational analysis. These regions provide candidate targets for in vitro screening of active compounds.

**Figure 9:**
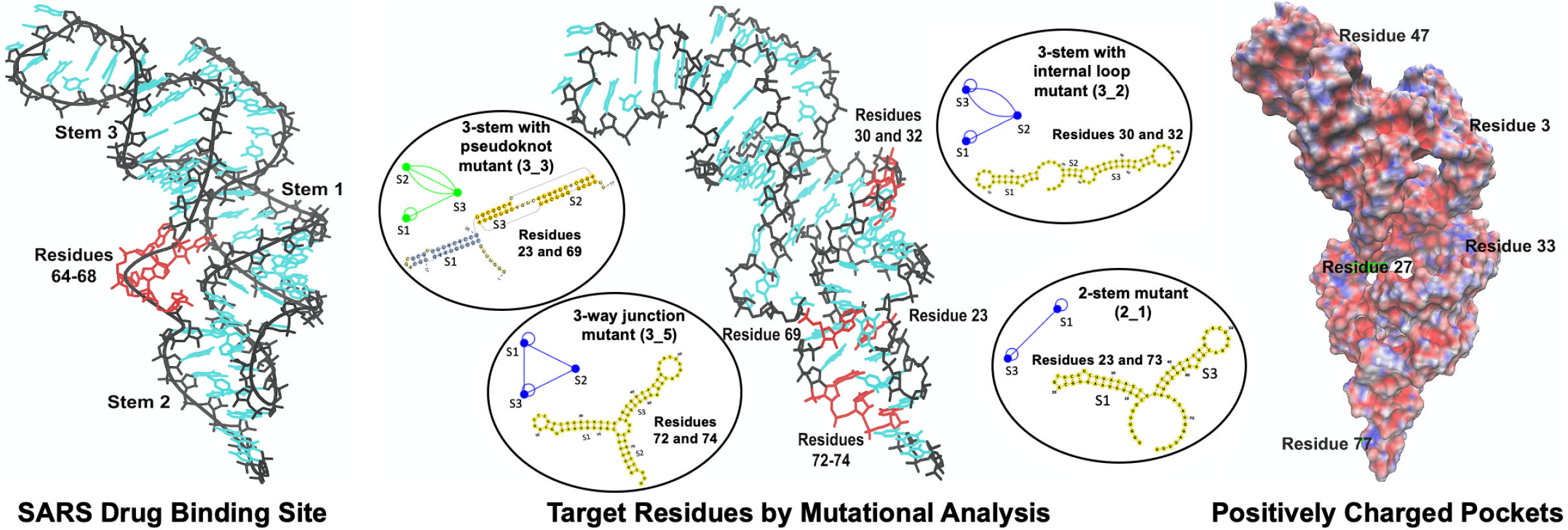
Target residues for drugs or gene editing emerging from this work. Resides correspond to previously identified drug-binding residues (10, 13), minimal mutations, and positively charged pockets that arose of our electrostatic analysis are shown for the frame-shifting element of the SARS-CoV-2 viral genome.

## 4 CONCLUSION AND DISCUSSION

We have proposed by computational design motif-altering minimal mutations of the SARS-CoV-2 FSE. In the key mutations highlighted, *only two residue changes* can accomplish a dramatic change in structure, namely destruction of Stem 2 and the pseudoknot. That the graph motif of the wildtype FSE can be recovered by compensatory mutations using our reverse design, in some cases with only one residue change, is also promising. Our MD simulations suggest overall stability of the wildtype and mutant system 2D structures. The tools developed here to design *in silico* RNAs containing pseudoknots are widely applicable to other design problems. The genetic algorithm in RAF-IF for inverse folding is very successful at generating a large number of sequences quickly and sorting out the sequence pools to obtain minimal mutations.

Besides these computational advances, the results produced here immediately suggest key residues as drug, as well as gene editing, targets for Covid-19 therapy. Already, for the SARS FSE, the small drug compound *1,4-diazepane derivative 10* (MTDB) has been shown to inhibit this frame-shifting process (10, 13). Recent *in vitro* MTDB binding experiments to the SARS-CoV-2 FSE demonstrated a similar inhibition effect (14). This lead, together with our identified mutation sites, as well as positively-charged regions of the RNA identified by electrostatic surface analysis of the FSE, are shown in Fig. 9. Underway are *in silico* screening for anti-viral compounds targeting the FSE with these residues. Viral-RNA based therapies that utilize CRISPR-like systems delivered through small-molecule vectors are also becoming more feasible.

Recently, a complete 2D structure prediction of SARS-CoV-2 genome was reported using *in vivo* SHAPE-MaP data (52). Two different FSE structures were predicted: one consistent with our 3-stem structure, and one forming a different Stem 3 with downstream sequence. The two possibilities were considered commensurate with the conformational flexibility of the pseudoknot. More thorough experiments are needed to confirm the FSE structure and examine its conformational flexibility.

As is well appreciated, various secondary-structure prediction programs use different algorithms and often produce different predictions. For the 77-residue system, 5 of the 8 prediction programs yield the 3-stem pseudoknot structure (Fig. 1). However, when the slippery site is added, only 2 of the 5 programs preserve the pseudoknot structure. Larger systems yield more diverse structures. The computational time for the different programs also varies significantly. For example, PKNOTS takes approximately 30 seconds for predicting the structure of 77 nucleotides, but with a time complexity *O*(*n*^6^), where *n* is the number of residues, large systems are too costly to handle. Programs like IPknot have a more favorite size scaling, and replacing PKNOTS with IPknot in our RAG-IF pipeline will allow us to extend our design pipeline to longer RNAs. The use of our graph-based tools is critical here for mutational transformations, as it provides us a robust coarse-grained approach insensitive to small differences in 2D structures.

Clearly, the overall fold of the FSE may depend on neighboring residues and on complexes with the nucleocapsid protein in the cellular context. In Fig. 10, we show predicted structures for the wildtype and 3-way junction mutant 3_5 in the context of different flanking sequences. The differences as a function of sequence length that emerge also reflect differences between programs. Interestingly, in some cases, a large sequence can recover our 77-nt consensus structure better than a shorter construct. Thus *in vivo* imaging and other experiments like SHAPE reactivity probing (54) would be important for understanding the contextual folding of this important region of the virus explored in this work in isolation.

**Figure 10:**
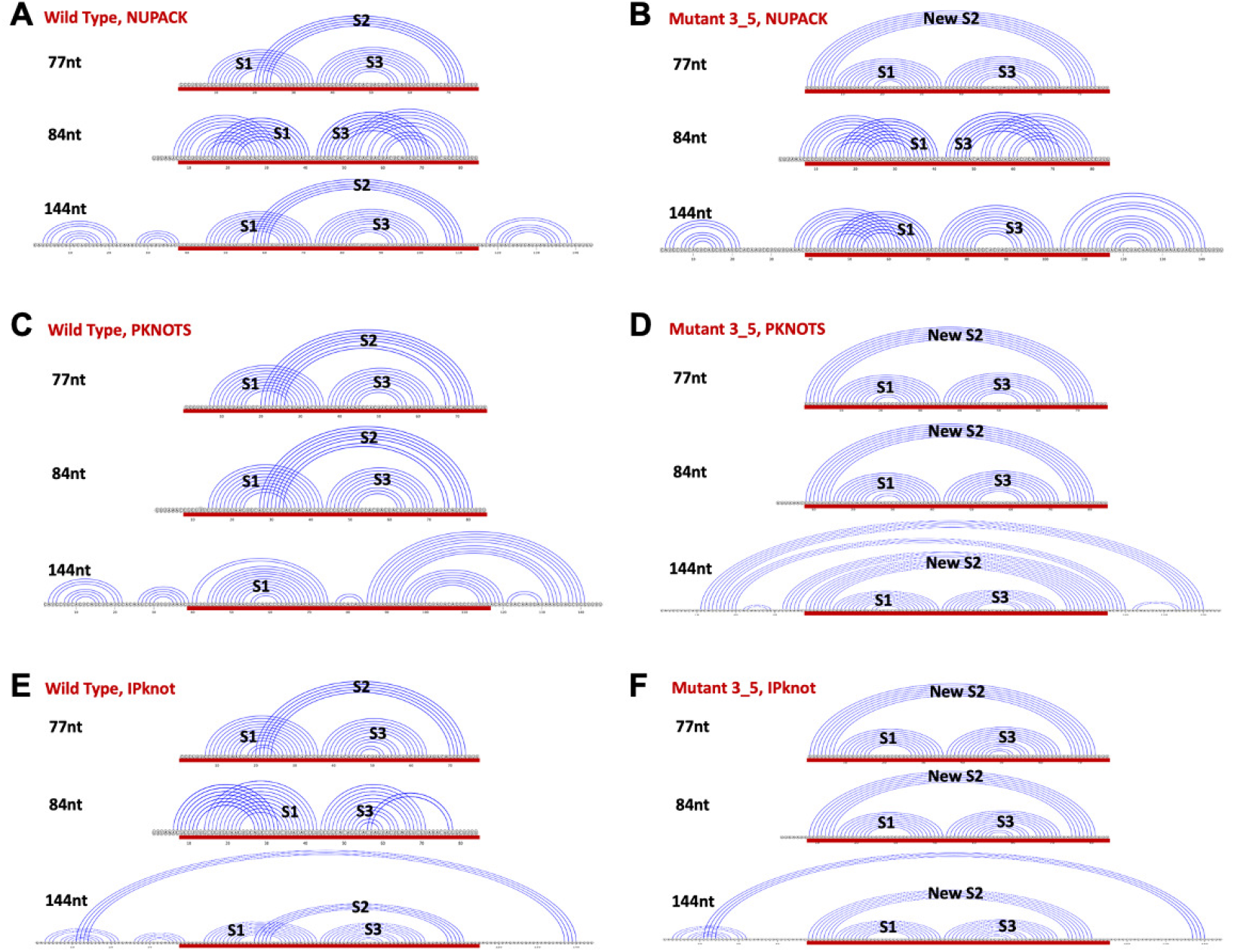
Predicted structures for the wildtype and mutant 3_5 using different sequence lengths. The 84nt sequence contains the slippery site, and the 144nt sequence adds 30nt to both ends of the 84nt sequence. The sequences (aligned with common 77nt subsequence, red) are shown for: (A) wildtype FSE/NUPACK (30), (B) 3-way junction mutant 3_5/NUPACK, (C) wild-type/PKNOTS (29), (D) 3_5/PKNOTS, (E) wild-type/IPknot (31), (F) 3_5/IP-knot.

Because accumulating data show that SARS-CoV-2 is actively acquiring new mutations that allow it to escape known protein-targeting drugs, alternative therapeutic strategies which target the RNA viral genome, rather than viral shell or auxiliary proteins, present a flexible and efficient approach to counter new infections. By examining the 73,739 aligned viral sequences available in the GISAID database (53) as of August 10, 2020, in Fig. 6B, we found that only 1316 sequences (1.78%) have mutations in the 84nt FSE region. Among these, 1268 contain single point mutation, underscoring the high conservation of the FSE region. These mutations are shown along with the residues we identified for top minimal mutations in Fig. 6. Our mutation residues 23 and 74 are not among current variants, but residues 69, 72 and 73 occur with one mutation to U each; residue 32 is mutated twice to G, and residue 30 is mutated 7 times to G. This generally high sequence conservation of the FSE and stability to mutations suggests it is a good drug target.

Indeed, drug binding of MTDB to six FSE natural point mutations suggested insensitivity to natural mutations (55). The mutational transformations we reported here provide insights for further *in silico* and ultimately *in vitro* drug screening to find other therapeutically effective drugs. In general, understanding the structure of the FSE and other gene regions of coronaviruses is important for helping address future viral threats which inevitably may arise. Such knowledge and methodologies are also transferable to other coronaviruses or other viruses that could pose threats to human health in the future.

Further SARS-CoV-2 viral genome variant analyses are also important for drug discovery and for identifying the spread of Covid-19 around the globe. Initially, it was believed that different strains are not related to virulence or infectivity, but theories are evolving, as associations of different genomic mutations with different levels of infectivity are being debated (e.g., (56)). As more data on associations of viral mutants and disease outcome accumulate, further analysis on the viral genome will be invaluable.

In closing, molecular modeling and computational experiments such as described here have an important role to play in this fight against the small RNA enemy which has upended our modern world as never before since World World II. The continued interaction and participation of the biophysics, high-performance computing, and bioinformatics/computational biology communities will be invaluable in this quest. An improved understanding of the complex structural biophysics of the RNA viral genome will undoubtedly be useful and relevant to other coronaviruses, whose potential to emerge in the future cannot be understated.

## 5 AUTHOR CONTRIBUTIONS

TS, QZ, SJ, and SY designed the research. TS, QZ, SJ, and SY performed the computational study and analyzed the results. TS, QZ, SJ, and SY wrote the article.

## 6 ACKNOWLEDGMENTS

We gratefully acknowledge funding from the National Science Foundation RAPID Award 2030377 from the Division of Mathematical Science and the Division of Chemistry; National Institutes of Health R35GM122562 Award from the National Institute of General Medical Sciences; and Philip-Morris International to T. Schlick. We thank Shereef Elmetwaly for technical assistance, and David Ackerman, Stratos Efstathiadis, and Shenglong Wang from the NYU High Performance Computing facilities for providing our group dedicated resources to perform this work.

